# The fruit fly *Anastrepha obliqua* harbors three kingdoms of life in its intestinal tract

**DOI:** 10.1101/2021.07.06.451314

**Authors:** GR Amores, G Zepeda-Ramos, LV García-Fajardo, Hernández Emilio, Guillén-Navarro Karina

## Abstract

The fruit fly *Anastrepha obliqua* is an economically important pest for mango fruits in Mexico. The sterile insect technique is used to control this pest; it involves mass production and release of sterile flies to reduce reproduction of the wild population. As noted in different tephritidae, the performance of sterile males may be affected by the assimilation of nutrients under mass-rearing conditions. In the wild, the fly’s life cycle suggests the acquisition of different organisms that could modulate fitness and physiology of the fly. Therefore, the microorganisms lodged in the gut may be determinative. For *A. obliqua*, there is no information regarding microorganisms other than bacteria. This study analyzed bacteria, fungi, and archaea communities in the *A. obliqua* gut through denaturing gradient gel electrophoresis (DGGE) profile of 16S and 18S ribosomal DNA markers. Besides, 16S sequencing and phylogenetic analysis provided a better description of bacteria, and archaea communities. We found that wild flies presented higher microbial diversity than laboratory samples. Phylogeny analyses of wild samples suggest the presence of microbial species related to fructose assimilation while laboratory microbial species suggest the presence of microorganisms leading to a specialized metabolism to process yeast as result of the consumption of an artificial diet. Here, the archaea kingdom is suggested as an important player in fly metabolism. This is the first report of the intestinal microbial (bacteria, archaea and fungi) composition of *A. obliqua*, which will aid in our understanding of the role of microorganisms in the development and physiology of the flies.

## Introduction

Fruit flies (Diptera: Tephritidae) encompass ~70 species (Jurkevitch, 2011), which infect more than 30 fruit species, leading to worldwide economic impacts (Qin *et al*., 2015). In Mexico, despite being cosmopolitan, the Mediterranean fruit fly *Ceratitis capitata* (Szyniszewska and Tatem, 2014) has been controlled by the sterile insect technique (SIT). However, the fruit fly *Anastrepha obliqua* (Macquart) is a pest that causes losses of fruit crops; from January to July 2018 there were losses of 22 million tons of mango (*Mangifera indica*) and spondias (*Spondias purpurea* and *S. mombin*) fruits (SENASICA, 2018). SIT is a biological technique without adverse impact on biodiversity and the environment. SIT involves the systematic mass release of irradiated sterile adult competitive and flying males, which induce sterility in the wild population, by preventing offspring (KNIPLING, 1959; Montoya P. Toledo J., 2010).

Different efforts had been made to optimize the efficiency of SIT by enhancing the quality of sterile insects(Shelly and McInnis, 2001; Ami, Yuval and Jurkevitch, 2010). In this sense, nutrient assimilation is a critical factor for the quality of the released sterile insects. Composition of the artificial diet, enzymatic metabolic machinery, and microorganisms harbored in the fly’s gut modulate assimilation, contributing to fly fitness (Nestel, Nemny-Lavy and Chang, 2004; Rivera-Ciprian *et al*., 2017; Rempoulakis *et al*., 2018). Additionally, the wildlife cycle of the fruit fly suggests the acquisition of different microorganisms, which exploit nutrients from the natural diet, modulating its biology (Domínguez J. Artiaga-López T., 2010; Ben-Yosef *et al*., 2014). Therefore, the fly’s gut ecology must be determinative in the modulation of fly fitness.

The microorganisms present within a host, known as microbiota, are generally well known for modulating host health and fitness (Sommer *et al*., 2016; Thaiss *et al*., 2016). In fruit flies, most of the microbial diversity studies have focused on elucidating the role of bacteria housed in the gut of different fly genera, such as *Ceratitis*, *Drosophila*, *Bactrocera*, *Helaeomyia*, and *Anastrepha* (Kadavy *et al*., 2000; Kuzina *et al*., 2001; Capuzzo *et al*., 2005; Juneja and Lazzaro, 2009; Ben-Yosef *et al*., 2014). Regarding these, bacterial microbiota such as *Klebsiella oxytoca*, *K. pneumoniae*, *Citrobacter freundii*, *Enterobacter* sp. and *Providencia rettgeri* have been evaluated as probiotics to enhance fly fitness (Ami, Yuval and Jurkevitch, 2010; Hamden *et al*., 2013; Augustinos *et al*., 2015; Roque-Romero *et al*., 2020). Therefore, manipulation of resident bacterial populations could have an effect on host fitness, possibly by generating stronger sterile males, which will compete better against wild flies(Ami, Yuval and Jurkevitch, 2010; Kyritsis *et al*., 2017). In addition, fungi and actinomycetes studies have focused on evaluating their role as insecticides, leading to the proposed use of different strains, such as *Beauveria bassiana*, *Metarhizium anisopliae*, and *Bacillus cereus* to improve biocontrol of *C. capitata* (Imoulan and Elmeziane, 2014; Navarro-Llopis *et al*., 2015; Ruiu *et al*., 2015; Samri *et al*., 2017). Interestingly, for the same fly species, different microorganisms have been proposed to present a functional role, indicating the importance of elucidating the gut ecology of the fly.

For the related fly, *Anastrepha ludens*, bacteria such as *Citrobacter*, *Enterobacter*, *Klebsiella*, *Providencia*, and *Pseudomonas* have been found in its intestinal tract (Kuzina *et al*., 2001). To date, studies on *Anastrepha* spp. have mainly focused on bacteria, and have not investigated other associated microorganisms, which could also modulate physiology and fitness. Thus, knowing the ecology of the fly intestine would provide a better understanding of microbiota–insect interactions, multi-species community structure, and its effect on the host. To determine the microbiota diversity associated with the gut of *A. obliqua*, we employed 16S and 18S PCR-denaturing gradient gel electrophoresis (DGGE) to describe the diversity of fungi, bacteria, and archaea present in the intestinal tract of wild and laboratory flies. Moreover, we used cloning techniques of 16S-ribosomal DNA (rDNA) coupled with phylogenetic analyses to gain better insight of the species harbored in the fly intestine. This knowledge will provide the basis for further studies of the microbiota in the digestive tract of *A. obliqua*, which could improve the success of sterile insect release.

## Materials and Methods

### Biological material

The larvae and adult *A. obliqua* flies were obtained from a mass-rearing colony at Moscafrut Facility in Metapa de Domínguez, Chiapas, Mexico. Third instar larvae were harvested and fed a diet containing 19% corn cob powder (Mafornu, Cd. Guzmán, Jalisco, Mexico), 5.3% corn flour (Maíz Industrializado del Sureste, Arriaga, Chiapas, Mexico), 7% torula yeast (Lake States, Div. Rhinelander Paper, Rhinelander, WI, USA), 9.2% sugar (Ingenio Huixtla, Chiapas, Mexico), 0.4% sodium benzoate (Cia. Universal de Industrias, S.A. de C.V., Mexico), 0.2% nipagin (Mallinckrodt Specialty, Chemicals Co., St. Louis, MO, USA), and 0.44% citric acid (Anhidro Acidulantes FNEUM, Mexana S. A. de C.V., Morelos, Mexico) (Orozco-Dávila *et al*., 2017). Lab adult specimens were collected after emerging from the pupae stage on vermiculite incubated at approximately 14 days and fed with hydrolyzed protein:sugar (in a 1:3 ratio), and water until collection at 12 days of age. Wild larvae were obtained from infested mango fruits provided by the Junta Local de Sanidad Vegetal del Soconusco, Chiapas. Third instars were obtained from infested mango fruits collected in the surroundings of the town of Tuxtla Chico, Chiapas, Mexico. Wild adult flies were caught at the end of 2010 harvest and during the 2011 harvest in mango orchards in the same town using modified MacPhail traps baited with hydrolyzed protein as an attractant to catch live adults (Enkerlin, 2018). Flies were taxonomically identified using the keys previously described (FAO/OIEA, 2018). All samples were washed twice with 70% ethanol. Dissections were performed to extract the digestive tract under sterile conditions in a laminar flow hood and stored in 70% ethanol at −20°C until processing.

### DNA extraction

Before extraction, samples (0.3 g) of the midgut were washed twice with 700 μl of 20 mM EDTA. DNA was extracted following the phenol–chloroform protocol (Sambrook, 2001) with some modifications. Briefly, 500 μl of 2 mM EDTA pH 8, 500 μl phenol pH 8.0, 0.25 g of glass beads, and 60 μl 20% SDS were added to macerated midguts and homogenized for 4 min. DNA purification was performed with chloroform and centrifuged at 13,000 x *g*. Then, 2.7 M sodium acetate and two equal volumes of absolute cold ethanol were added to the aqueous phase, which was incubated for 40 min at −70°C to precipitate DNA, centrifuged 35 min at 16,000 x *g*, washed with 70% ethanol, decanted, and allowed to dry at 37°C. The DNA pellet was resuspended in 50 μl water with 1.5 μg/μl RNAse and incubated for 1 h at 37°C. The obtained genomic DNA was purified using the Soil Microbe DNA kit MiniPrep ZR^™^ (Zymo Research, USA) according to the manufacturer’s instructions. Each sample was analyzed by 0.8% agarose gel electrophoresis to verify quality and purity.

DNA extraction from legs was performed using a single laboratory adult specimen that was frozen at −20°C. The legs were dissected, placed in a 1.5 ml tube, and washed with 70% ethanol five times. DNA extraction was performed as described above.

### PCR-DGGE analysis

Amplification of bacterial 16S rDNA fragments was performed with primers F984GC (CGCCCGGGGCGCGCCCCGGGCGGGGCGGGGGCACGGGGGGGCGCAACGCGAAGAACCTTAC) and R1378 (CGGTGTGTACAAGGCCCGGGAACG) (Nübel *et al*., 1996). Amplification of Actinomycetes 16S rDNA was performed with a nested PCR employing the Primer F243 (GGATGAGCCCGCGGCCTA) and R1378 (same as used in bacteria) for the first reaction to Actinomycetes population enrichment, and F984GC and R1378 in the second (Heuer *et al*., 1997).

To verify the presence of fungi, amplicons were obtained with primers NS1 (GTAGTCATATGCTTGTCTC) (White T. A. *et al*., 1990) and GCfung (GCCCGCCGCGCCCCGCGCCCGGCCCGCCGCCCCCGCCCCATTCCCCGTTACCG) (May, Smiley and Schmidt, 2001).

Archaea identification was performed using a nested amplification using primers 21F (TTCCGGTTGATCCYGCCGGA) and 958R (CCGGCGTTGAMTCCAATT) (DeLong, 1992) in the first PCR amplification; Parch519F (CAGCCGCCGCGGTAA) and Arch915R (CGCCCGCCGCGCCCCGCGCCCGGCCCGCCGCCCCCGCCCCGTGCTCCCCCGCCAATTCCT) (Coolen *et al*., 2004)were used for the second. One primer for each pair was designed with a GC clamp for DGGE analysis. The reaction mixture contained 0.20 ρmol of each primer, 1x buffer, 0.20 mM dNTPs (dinucleotide triphosphate), 1.5 mM MgCl_2_ (2.0 mM for fungi), 1 U *Taq* DNA polymerase (Invitrogen, USA), and template DNA for a total volume of 20 μl. Negative control included ultrapure water, whereas *Escherichia coli* DH5-α, *Streptomyces* sp., and *Trichoderma* sp., DNA were used as positive controls for bacteria, actinomycete and fungi, respectively. Amplification protocol in bacteria and actinomycetes consisted in a denaturation at 94 °C for 5 min, followed by 30 cycles of 1 min at 95 °C, 1 min at 58 °C and 40 sec at 72 °C, with a final extension at 72 °C for 5 min. The actinomycetes nested amplification used the same protocol for second amplification; for the first annealing temperature was modified at 53 °C (Das, Royer and Leff, 2007). To confirm the absence of actinomycetes in PCR negative samples, we determined the minimum amount of *Streptomyces* DNA that renders an amplicon using the corresponding primers. This PCR was done using serial dilutions (14 ng to 0.0014 ng) of DNA. Fungi PCR conditions consisted of a denaturation at 95 °C for 2 min, followed by 35 cycles of 30 sec at 95 °C, 30 sec at 50 °C, and 1 min at 72 °C, with a final extension at 72 °C for 5 min (Nikolcheva, Cockshutt and Bärlocher, 2003).

Archaea nested PCR protocol for the first reaction was: denaturation at 94 °C for 5 min, followed by 30 cycles of 40 sec at 94 °C, 40 sec at 53 °C, 1 min at 72 °C, and a final extension at 72 °C for 5 min. For the second reaction, alignment temperature was changed to 57 °C. Total DNA from an anaerobic plant sludge wastewater treatment was used as PCR positive control.

For DGGE, PCR products were quantified using Molecular Kodak Imaging software, and loaded in equal concentrations directly on a 0.8% polyacrylamide gel with a 20–50% denaturing gradient of urea and formamide, and electrophoresed in 1% TAE buffer (40 mM TAE, 2 mM Tris-acetate, and 1 mM Na_2_EDTA, pH 8.5). Electrophoresis was performed in the D-Code^™^ Universal Mutation Detection System (BioRad, Hercules, CA, USA) at 90 V for 8.5 h, with a constant temperature of 60°C. Subsequently, the gels were stained with SYBR Gold. Each DGGE was repeated three times in each experiment.

The DGGE band profiles indicate the different microbial communities from the fly midgut (Online Resource 2). The profiles were compared using the Jaccard index (J), which allows analysis of the biodiversity found in a sample where the maximum value indicates a greater diversity. This was calculated according to the following formula: J = nAB (nA + nB - nAB) – 1, where nAB is the number of bands in common between lanes A and B, nA is the total number of bands in lane A, and nB is the total number of bands in lane B.

The Shannon index (H’) allows comparisons of community similarity between two samples, where a value of 1.0 (or 100%) corresponds to communities that share an identical pattern, and a value of 0 indicates that no difference exists. This index was calculated using the following equation: H’ = - (ni/N) (log ni/N), where H’ is the diversity, and ni/N is the number of individuals of the species given by band intensity (ni) to the total subjects (N, the total intensity of all bands of the same sample); *i.e*., the relative abundance of species. Band intensity reflects the abundance of the same (Eichner *et al*., 1999) and theoretically represents a genus, in this case, analyzed by Molecular Kodak Imaging software.

### 16S cloning and phylogenetic analysis

The 16S rDNA amplicons for actinomycetes, bacteria, and archaea were obtained as described above. The purified PCR products were cloned into a p-JET vector with CloneJet PCR Cloning Kit (Thermo Scientific, Waltham, MA, USA) and transformed into *E. coli* DH5-α cells (Sambrook, 2001). Clones were sequenced using pJET primers by capillary sequencing at Macrogen Inc. (Korea). The sequence chromatograms were visually inspected and manually trimmed using AliView software (Larsson, 2014). To identify closely related 16S rRNA genes, the remaining sequences were analyzed with the BlastN tool at 99% identity against the 16S archaea, actinomycete, and bacteria database at NCBI as described in the main text. NCBI accession number of the sequences employed for phylogenetic analyses are shown at Online Resource 1. Multiple sequence alignment was performed using ClustalW (Thompson, Higgins and Gibson, 1994). The best suitable model and phylogenetic trees were constructed using Molecular Evolutionary Genetics Analysis (MEGAX) software and several algorithms (Kimura, 1980; Felsenstein, 1985; Tamura, 1992; Tamura *et al*., 2013). The accuracy of the tree topology was performed by 1000 bootstrap replicates (Felsenstein, 1985).

### Geneious diversity analysis

The 16S rDNA clones of bacteria and archaea communities obtained from wild and laboratory adult and larvae flies were loaded into Geneious software version 11.1 (Kearse *et al*., 2012). Subsequently, all submitted NCBI sequences as well as those which display robust electropherogram (Online Resource 3) were manually curated and merged to gain a general view of the global biological diversity harbored at *A. obliqua*. Clean sequences were then analyzed using the 16S diversity tool in Geneious software.

## Results and Discussion

### Bacteria communities

The microbiota interacts with its host, modulating nutrient assimilation, health and fitness. In the related fruit-fly pests, *Anastrepha obliqua*, *A. ludens*, and *Ceratitis capitata*, is known that gut enzymatic activity (Rivera-Ciprian *et al*., 2017) as well as bacteria regulates their food bio-assimilation capabilities producing fitness effects (Kuzina *et al*., 2001; Ami, Yuval and Jurkevitch, 2010; Gallo-Franco & Toro-Perea, 2020). Because of this, we got interested in disclosing *A. obliqua*’s bacterial microbiota diversity with an additional systematic search for actinomycetes due to its host-related biochemical metabolism. For this, DNA was extracted from the digestive tract of wild adult and larvae (AW and LW, respectively) and laboratory (AL and LL, respectively) flies, and samples were subjected to 16S rDNA bacterial and actinomycete PCR amplification (as described in methodology and further referred as Gram-positive population for actinomycete) followed by DGGE. We observe different patterns of migration between developmental stages and groups suggesting different bacterial communities. Besides, our DGGE Gram-positive enriched experiments over all samples recovered just three communities in the midgut of wild larvae flies sample. To confirm this, we tested a minimum amount of DNA for the absence of actinomycetes in other stages of the fly. We could only detect actinomycetes up to a minimum concentration of 0.14 ng of DNA, suggesting that if there are actinomycetes, they are found in undetectable amounts in the samples analyzed by DGGE as has been reported for taxa of low abundance (González *et al*., 2011). It has been demonstrated that the detection limit of PCR-DGGE is 104–108 CFU/ml (Ercolini, 2004), suggesting that the communities in our samples are below these limits or the primers were inefficient for actinomycetes in this fly. Indeed, amplification of 16S rDNA for actinomycetes showed smear bands (data not shown) in all fly stages, which were subsequently used for cloning.

In general, population index analyses show that adult flies harbor higher diversity than larvae. Specifically, Shannon index (H’) analyses (Table 1) showed that adult wild flies presented the greatest bacterial diversity (1.22 H’), as expected in a native environment with a varied diet, which could help establish a more diverse group of microorganisms. The wild larvae showed inferior diversity (1.07 H’) than the adult, as its development is limited within the host fruit. Furthermore, Jaccard index (J) analysis (Table 1) showed that LW and AW share 53% of their bacterial communities, suggesting the same vertical transmission that has been observed in *C. capitata* (Behar, Yuval and Jurkevitch, 2008), where the female establishes populations of microorganisms during oviposition, which may contribute to optimal development in the larval stage and some are kept during adulthood. Regarding mass-rearing flies, larvae and adult samples showed lower diversity (H’ = 0.94 and 0.95, respectively) than wild flies and shared 63% of their communities. This could be due to a similar diet and confined environment. It is worth mentioning that, contrary to other works, the adult wild flies were collected as adults from mango orchards and not from fruits infested with larvae letting remain in lab conditions until emergence, so wild gut microbiome was better represented. Comparison of mass-rearing and wild flies showed that larvae samples share 52%, while adult flies share only 36% of their microorganism communities. Since the mass-rearing *A. obliqua* colony has been maintained for more than 150 generations, this allows us to suggest that it must contain bacterial core microbiota from both natural and mass-rearing environments, which could contribute to metabolic function of the flies.

**Table 1.** Diversity (H’) and Similitud (J) indexes of the microbial communities found in the midgut of A. obliqua

| Sample | Bacteria |  |  | Archaea |  |  | Fungi |  |
| --- | --- | --- | --- | --- | --- | --- | --- | --- |
|  | H | J % | J %* | H | J % | J %* | H | J % |
| **AW** | 1.22 | 53<sup>1</sup> | 36 | 1.0 | 77 | 78 | 0.8 | 62 |
| **LW** | 1.07 |  | 52 | 0.77 |  | 85 | 0.67 |  |
| **AL** | 0.95 | 63<sup>2</sup> | 36 | 0.8 | 83 | 78 | 0 | 0 |
| **LL** | 0.94 |  | 52 | 0.77 |  | 85 | 0 |  |

Shannon (H') and Jaccard (J) indexes were determined from DGGE as described in the methodology. Adult wild (AW), Larvae wild (LW), Adult laboratory (AL) and Larvae laboratory (LL). Comparison between AW and LW (%<sup>1</sup>); Comparison between AL and LL (%<sup>2</sup>); Comparisons between adults and between larvae samples independently of their environmental origin (%\*) (i.e. AW-AL; LW-LL).

Upon bacteria diversity evaluation, we were interested in identifying the species that could be harbored in the fly midgut. With this in mind, amplified and cloned specific 16S rRNA gene region sequences were blasted against the 16S bacteria NCBI database, and a phylogenetic tree was constructed. We obtained 30 bacterial and 23 actinomycetes sequenced clones; after analysis, we kept 15 bacteria and 3 actinomycetes unique sequences. Thus, taxonomic classification of these sequences was restricted because of the low throughput cloning and sequencing methods.

The phylogenetic analysis of bacteria cloned sequences showed two main groups (Fig. 1) composed of Gram positive and negative bacteria. In general, *Enterobacteriaceae* is the most common group found in the intestinal tract of fruit flies (Kuzina *et al*., 2001; Behar, Yuval and Jurkevitch, 2005, 2008) as shown here for all samples. Our results showed that the majority of the mass-rearing larvae sequences were clustered within an Enterobacteriaceae group composed of *Providencia*, *Klebsiella*, *Serratia*, *Pectobacterium*, and *Morganella*, but were undetected in non-irradiated laboratory adult samples (Fig. 1 compare light-dark blue circles). In adult flies, microbiota composition is mainly studied in mass-rearing individuals with evidence of decreases on those Enterobacteriaceae strains, but maintaining populations of *Klebsiella*, *Providencia* and *Enterobacter*, by adding them as probiotics after irradiation improves mating capabilities and fly development (Ami, Yuval and Jurkevitch, 2010; Augustinos *et al*., 2015; Liu *et al*., 2016; Roque-Romero *et al*., 2020), supporting the importance of those strains in the metabolism of the fly. One LL sequence was related to *Aerococcus viridians* and *Enterococcus fecalis*. These bacteria species have been differentially described in wild and mass-rearing larvae and adult flies of *Drosophila melanogaster*, *Ceratitis capitata*, *Bactrocera* spp., and *Anastrepha ludens* (Juneja and Lazzaro, 2009; Ami, Yuval and Jurkevitch, 2010; Wang *et al*., 2014). And, those bacterial strains are related to the diet nutrients metabolisms in flies such as urea metabolization into amino acids, suggesting that these bacterial strains could contribute to nutrient assimilation (Ben-Yosef *et al*., 2014, 2015). The mass-rearing adult sequences clustered within metabolically specialized species of bacteria, such as members of the *Desulfovibrionaceae* family and *Lentisphaerae phylum* (i.e., *Victivallis* and *Oligosphaera* spp.) (Zoetendal *et al*., 2003; Fuerst, 2013; Qiu *et al*., 2013). Sulfate reducing bacteria (*Desulfovibrionaceae* family) are found in humans (Loubinoux *et al*., 2002), where they modulate the metabolism of primary fermenter bacteria in the intestine to regulate energy supply; they have also been linked to bowel disease (Wegmann *et al*., 2017). In insects, *Desulfovibrionaceae* family bacteria are found in the beetle *Amblonoxia palpalis* with no effect in their populations (Koneru *et al*., 2016). *Lentisphaerae*-isolated bacteria are related to sugar fermentation (Qiu *et al*., 2013). The presence of most of the adult sequences at this bacteria group suggest that adult laboratory flies develop a specialized microbiota related to mass-rearing process.

**Fig. 1.**
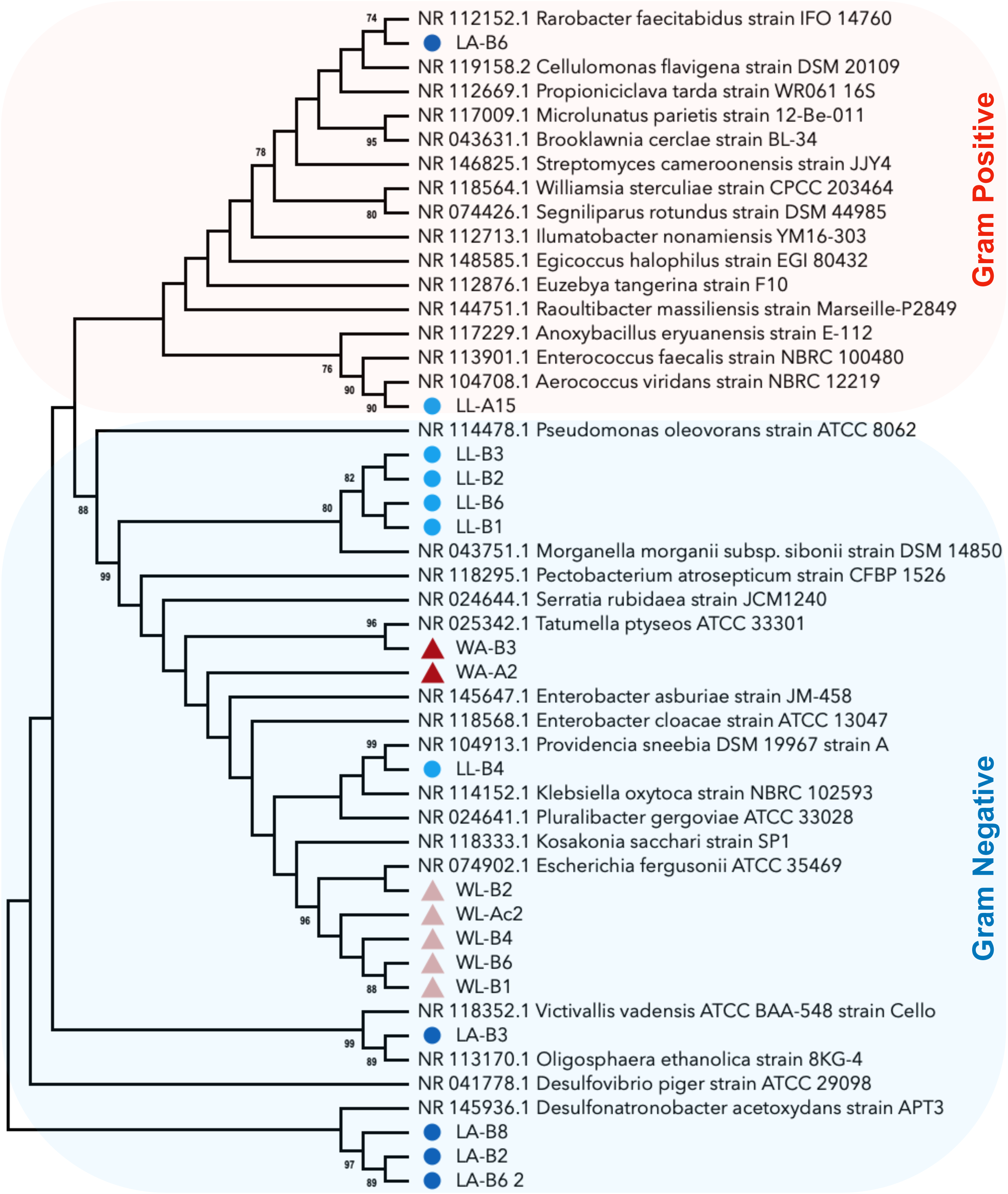
Phylogenetic tree of bacteria species identified in the intestine of *Anastrepha obliqua*. Gram negative and positive bacterial communities are delimited in blue and red, respectively. The 16S ribosomal DNA (rDNA) bacterial clones obtained from wild adult and larvae flies are shown as WL_ (pink triangles) and WA_ (red triangles); the laboratory samples as LL_ (blue light circles) and LA_ (blue circles). The tree (439 positions) was constructed using maximum composite likelihood (MCL) with a Kimura 2-parameter model. The percentage higher than 70% of 1000 bootstrap resampling is shown next to the branches. Evolutionary analyses were conducted using MEGAX.

In the Gram-positive bacterial group, one mass-rearing adult sequence was related to *Rarobacter* species. Isolated *Rarobacter* species can use a broad spectrum of carbohydrates, such as D-fructose and polyols, in the presence of proteinaceous substrates or inorganic nitrogen, and can also produce proteases to lyse yeast (Shimoi and Tadenuma, 1991). These microbial populations might play an important metabolic role within the fly, since artificial diets contain different compounds, such as yeast (Domínguez J. Artiaga-López T., 2010), but also contain preservants, such as methylparaben, which can affect microbial populations and mate choices, as has been shown in *D. melanogaster* (Obadia *et al*., 2018). Altogether, in addition to the irradiation process, the mass-rearing conditions and biological processes within the fly could also be responsible for controlling the bacteria load. These results suggest the importance of the microbial species described here in modulating the metabolism of mass-rearing *A. obliqua* fruit flies, although their functions during the nutrient assimilation process have yet to be determined. Our results also suggest that microbiota and host interactions help establish the proper metabolic conditions to modulate fly fitness.

Regarding wild samples (Fig. 1, triangles), larvae sequences were related to the *Enterobacteriaceae* (*Escherichia* and *Kosakonia*) and *Pseudomonadaceae* families. *Kosakonia sacchari* is a nitrogen-fixing bacterium reported in *Saccharum officinarum* L. However, the isolated strain uses sugars such as fructose and glucose (Chen *et al*., 2014); thus, this strain could metabolize sugar. *Pseudomonas* spp. has been identified in adult wild *C. capitata* and adult laboratory *A. ludens* (Kuzina *et al*., 2001; Behar, Yuval and Jurkevitch, 2008). Behar *et al*. showed in 2008 that *P. aeruginosa* regulates fly longevity by modulating the endemic *Enterobacteriaceae* population in the midgut of *C. capitata* flies. These strains are also present in artificial diets and induce beneficial effects during larval development and in adult flies (Augustinos *et al*., 2015; Rempoulakis *et al*., 2018). Moreover, the presence of strains such as *Pseudomonas*, *Erwinia*, and *Escherichia* also suggests that flies can be considered as reservoirs; in flies, strains of the same genus were found to induce disease in fruits(Kadavy *et al*., 2000; Sela *et al*., 2005; Kapsetaki *et al*., 2014; Ordax *et al*., 2015). The wild adult sequences were related to *Tatumella* and *Pluralibacter*, and *Pluralibacter* was described together with *Enterobacter cloacae* in *A. ludens* (Kuzina *et al*., 2001). Presence or addition as probiotics of those strains induce an enhanced fitness in mass-rearing sterile flies (Ami, Yuval and Jurkevitch, 2010), supporting its role in wild fitness success. Altogether, we present evidence of the Gram-positive and negative bacteria diversity harbored in the midgut of wild and mass-rearing *A. obliqua* flies. These results are consistent with the microbial communities reported in *Anastrepha* related flies (Ventura *et al*., 2018; Gallo-Franco and Toro-Perea, 2020; Roque-Romero *et al*., 2020). The identification of different species suggests molecular crosstalk between these microorganisms and the host, which is important in metabolic pathways such as carbon and nitrogen uptake. It remains to be determined whether those communities are involved in commensalism, parasitism, or symbiosis.

### Archaea communities

Archaea are distributed in extreme and moderate habitats in which they play a significant role because of their metabolic differences compared to prokaryotic and eukaryotic organisms (Hara *et al*., 2005; Pikuta, Hoover and Tang, 2007; Bräsen *et al*., 2014; Könneke *et al*., 2014). For insects such as cockroaches, termites, and beetles, which can survive on a low-nutrient diet, archaea of the class *Halobacteria* and *Methanomicrobia* provide the proper conditions in the intestine (e.g., CH_4_) for the development of other bacteria that modulate host metabolism (Gijzen *et al*., 1991; Donovan *et al*., 2004; Ceja-Navarro *et al*., 2014). Due to its biochemical and biological importance, we were interested in analyzing the diversity of archaea communities in the gut of *A. obliqua*. The Shannon index showed that adult flies (wild, H’ = 1 and mass-rearing, H’ = 0.8) have greater diversity compared to larval stages (wild and mass-rearing, H’ = 0.77) (Table 1). The trend, as in bacterial communities, is that the wild adult exhibits greater diversity than wild larvae and mass-rearing samples, possibly due to a greater variety in their diet. The Jaccard index showed that wild specimens have similar archaea communities (77%), whereas the adult and larvae mass-rearing flies share 83% of their communities.

Besides, a comparison between adult and larvae stages also showed that they share more than 70% of their communities. Specifically, the AW and AL samples share 78%, while LW and LL share 85% of their archaea communities. As stated, archaea are endowed with particular pathways to produce CH_4_, but also modify other metabolic pathways, such as the Embden-Meyerhof-Parnas and Entner-Doudoroff pathways, which are related to sugar fermentation (Bräsen *et al*., 2014); this could explain the presence of similar archaea communities between the individuals analyzed as a requirement for the proper metabolism of nutrients. Overall, we suggest that archaea may be essential throughout the cycle life of the fly, and likely could be determined by the mother and diet, as described and supported above in the case of bacteria.

To gain a better perspective of the archaea species present in the midgut of *A. obliqua*, we cloned the 16S PCR product prior to sequencing. We got 45 sequences and after NCBI submission we kept 29 sequences. These primers have been shown to detect bacterial strains (Coolen *et al*., 2004). Therefore, we blasted our sequences against the 16S archaea database and, for the sequences with no match; a subsequent blast against the 16S bacteria databases at NCBI was performed. The sequences were analyzed as described in the methodology and a phylogenetic tree was constructed. Two main clades were observed, archaea and bacteria (Fig. 2). Interestingly, bacterial species detected with archaea-directed primers were different from the species identified with bacteria primers (compare Fig. 1 with bacteria clade in 2).

**Fig. 2.**
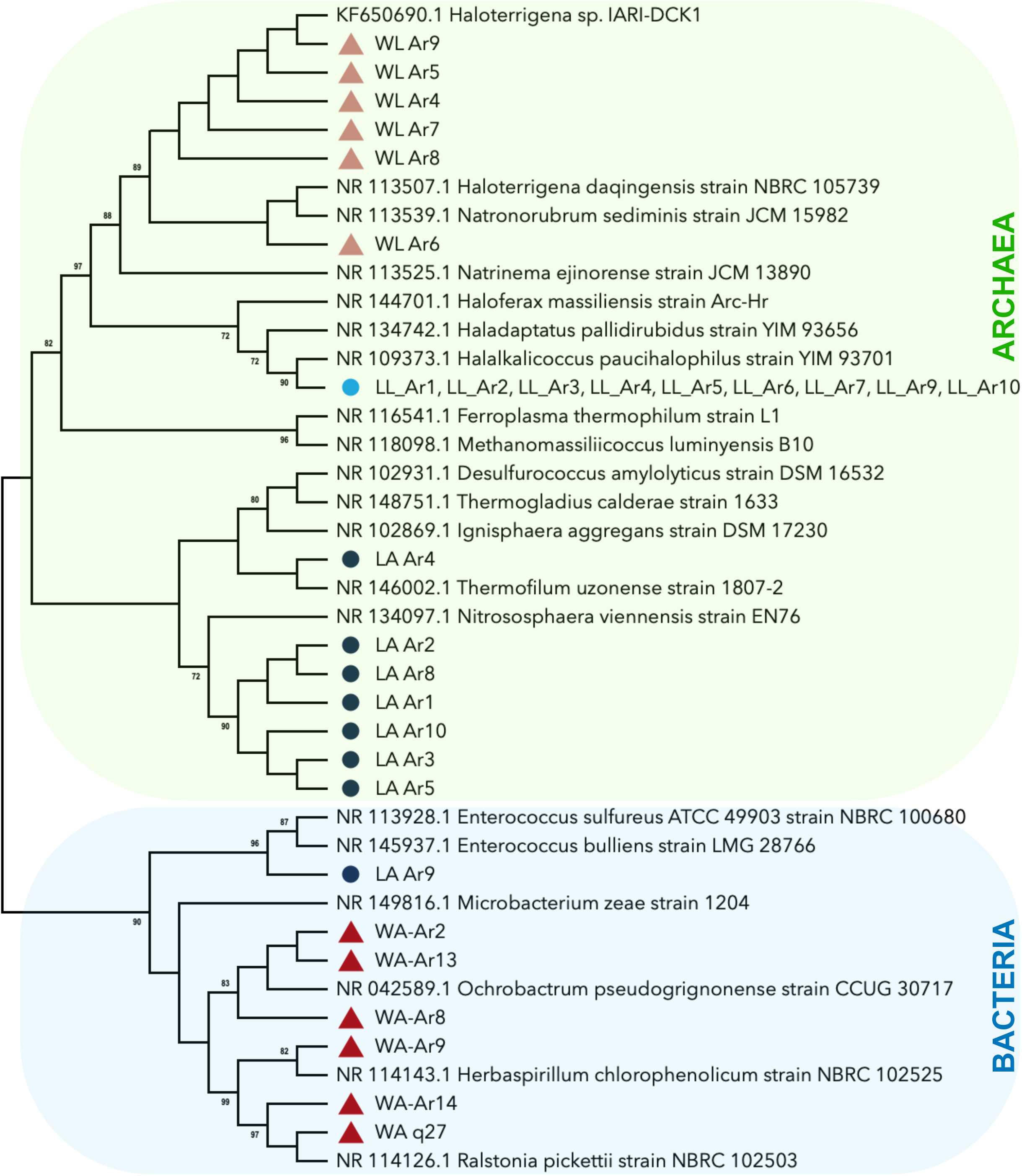
Phylogenetic tree of archaea communities identified in the intestine of *A. obliqua*. Two main clades can be seen, archaea (green) and bacterial (blue) communities. The 16S rDNA bacterial clones obtained from wild adult and larvae flies are shown as WL_ (pink triangles) and WA_ (red triangles); the laboratory samples as LL_ (blue light circles) and LA_ (blue circles). DAMBE software was employed to merge equal sequence clones. The neighbor-joining method to a matrix of pairwise distances estimation using the MCL approach with a Kimura 2-parameter model was used to construct the tree (406 positions). The percentage higher than 70% of 1000 bootstrap resampling of trees in which the associated taxa clustered together is shown next to the branches. Evolutionary analyses were conducted using MEGAX.

Mass-rearing samples were mainly clustered at the archaea clade and one AL sequence clustered at the bacteria clade within the *Enterococcus* branch, which is associated with lactic acid production (Kadri *et al*., 2015). Mass-rearing larvae were related to *Halalkalicoccus* species (*Halobacteriaceae family*), which can grow on fructose and mannose in the laboratory (Xue *et al*., 2005; Poehlein *et al*., 2016). Adult laboratory samples were related to strains that use inorganic compounds as an energy source but also organic compounds, such as yeast. These strains were *Nitrososphaera* spp., and *Thermofilum spp*., which can grow in the presence of ammonia, yeast, and peptone (Tourna *et al*., 2011; Toshchakov *et al*., 2015). Knowing the nutritional content of the artificial diet employed for mass-reared flies, and the microbial species found, our results suggest that those strains could be gained via the diet, modulating the bio-assimilation process since laboratory flies are reared with artificial diets enriched with yeast.

In the wild samples, larvae sequences clustered within the *Haloterrigena*, *Natronorubrum*, *Natrinema*, and *Haloferax* branch (*Halobacteriaceae* family). These species are related to phosphorus solubilization and can grow in the presence of fructose (Castillo *et al*., 2006; Yadav *et al*., 2015). Interestingly, wild larvae were collected from mangoes, and it is possible that these species could be enriched, as the mango has a high content of fructose and phosphorus. Therefore, these strains could be ingested with the natural diet, or acquired through vertical transmission, contributing to the host’s sugar assimilation. Finally, adult wild sequences clustered within the bacteria clade and no archaea was detected. Isolated *Ralstonia* and *Herbaspirillum* strains are associated with copper bio sequestration and degradation of chlorophenol, respectively (Im *et al*., 2004; Yang *et al*., 2010). Copper compounds are used in different crops, such as tomato, cucumber, and mango (Cazorla *et al*., 2002; İşeri *et al*., 2011). In mango crops, copper is used to avoid necrosis induced by *Pseudomonas syringae* pv. *Syringae* (Cazorla *et al*., 2002). In flies, depending upon the dose, this compound leads to positive and negative effects. Positively, copper works by activating superoxide dismutase to regulate oxidative stress, modulating the genotoxic effects of reactive oxygen species and therefore, fly survival increases; while copper intoxication leads to the opposite outcome (Marchal-Ségault, 1993; Matsuo, Ooe and Ishikawa, 1997; Arcaya *et al*., 2013; Southon, Burke and Camakaris, 2013; Carmona *et al*., 2015). This suggests that the species harbored in the *A. obliqua* intestine could modulate fly fitness by regulating copper and oxidative metabolism.

To the best of our knowledge, this is the first record of archaea in the digestive tract of *A. obliqua*. Archaea are important organisms for metabolism in the intestines of insects (Ceja-Navarro *et al*., 2014). It has also been described that archaea produce metabolites that are used by insects to locate fruits (Piñero *et al*., 2015). Thus, wild microorganisms could promote food pre-digestion, change the nutritional content, or improve digestion in the fly gut (Lemaitre and Miguel-Aliaga, 2013). Altogether, the results presented here suggest that archaea and bacteria strains, which could be ingested during natural or artificial feeding, could modulate metabolic pathways in tephritidae through the digestion and assimilation of nutrients. Therefore, the differences between archaea and bacteria specimens detected among our samples can highlight their importance in the midgut.

### Fungi diversity in wild and laboratory flies

Fungi provide sustenance and dwellings for insects, while insects offer material for fungal decomposition, protect growing spaces, and allow for transportation to new locations; these fungi-insect exchanges have been described as convergent interactions (Bittleston *et al*., 2016). In the intestine of *D. melanogaster*, 45 different species of fungi have been identified, and this diversity is related to the feeding environment (Stefanini, 2018). For the fruit fly *Ceratitis capitata*, isolated species of fungi, as well as their metabolites, have been suggested to act as biocontrol agents (Ortiz-Urquiza *et al*., 2009; Imoulan and Elmeziane, 2014; Navarro-Llopis *et al*., 2015; Ruiu *et al*., 2015). For *Bactrocera* spp., yeast of the genera *Hanseniaspora* and *Pichia* are fundamental in larval survival (Piper *et al*., 2017). Therefore, fungi must also play an essential role in the intestine of *A. obliqua*. Consequently, we evaluated the presence of fungi in the midguts of all samples. *A. obliqua* (laboratory adult) 18S-DNA amplicon from a leg (FL) was used as a control to discard bands that were also present in the FL pattern in DGGE analysis. This assay was repeated three times to ensure reproducibility.

Fungal communities observed in the intestine of wild adults had greater diversity than wild larvae samples. The wild adult fly exhibited greater diversity (H’ = 0.8) than wild larvae (H’ = 0.67), sharing 62% of their communities. In the case of the laboratory samples, a faint FL band pattern was observed with DGGE, restricting the H’ and J analyses (Table 1). We did not obtain sequenced clones because host DNA was always in greater quantity than the fungi it harbored. Nevertheless, our results suggest that fungal diversity in wild samples was significantly different compared with laboratory flies. Therefore, the fungal community likely exerts a considerable impact on the biology of *A. obliqua* either through a symbiotic relationship or as a pathogen; it is known that the fungal community can play essential roles in insect guts, and has been shown to produce toxins that are pathogenic for insects, such as beetles and black flies (Kostovcik *et al*., 2015; Varotto Boccazzi *et al*., 2017).

As beneficial guests, fungal communities could act as suppliers of organic ingredients, essential vitamins, and enzymes that promote digestion (Vega and Dowd, 2005). Fungi could also improve metabolism via sugar fermentation and nitrogen-fixing, as well as participate in pheromone production by synthesizing steroids, as has been shown in bark beetles (Klepzig *et al*., 2009). It is possible some of these functions occur in the digestive tract of wild larvae and adult flies, which could explain the diversities found between wild and laboratory samples.

### 16S rDNA hypervariable region diversity

Our study aimed to increase our knowledge regarding *A. obliqua* microbiota. Through this process, we found that there are some biases that must be overcome to determine microbial diversity with the best possible accuracy. These biases start with the selection of DNA extraction methods, primer design, and PCR yield (Guillén-Navarro *et al*., 2015). We showed that with the use of different primer sets aimed at the same target (diverse hypervariable regions of bacteria and actinomycetes 16S rDNA), it was possible to identify species that have not been previously described in this fruit fly. Next generation sequencing methods using 16S rDNA would allow us to increase the number of species identified, but could introduce biases due to different factors, one of which is the use of primers targeted to only one or two 16S rDNA hypervariable regions, which limits species identification in complex bacterial populations (Laursen, Dalgaard and Bahl, 2017). Thus, the phyla *Proteobacteria*, *Actinobacteria*, and *Deinococcus* have been favored as the most abundant in *Anastrepha ludens*, *A. obliqua*, *A. serpentina*, and *A. striata* through 16S pyrosequencing, with primers targeted to the V3 hypervariable region (Ventura *et al*., 2018). Our results also broaden our knowledge of the microbiota of *A. obliqua* with a 28% global abundance of the Archaea kingdom (Fig. 3 and Online Resource 4, 5).

**Fig. 3.**
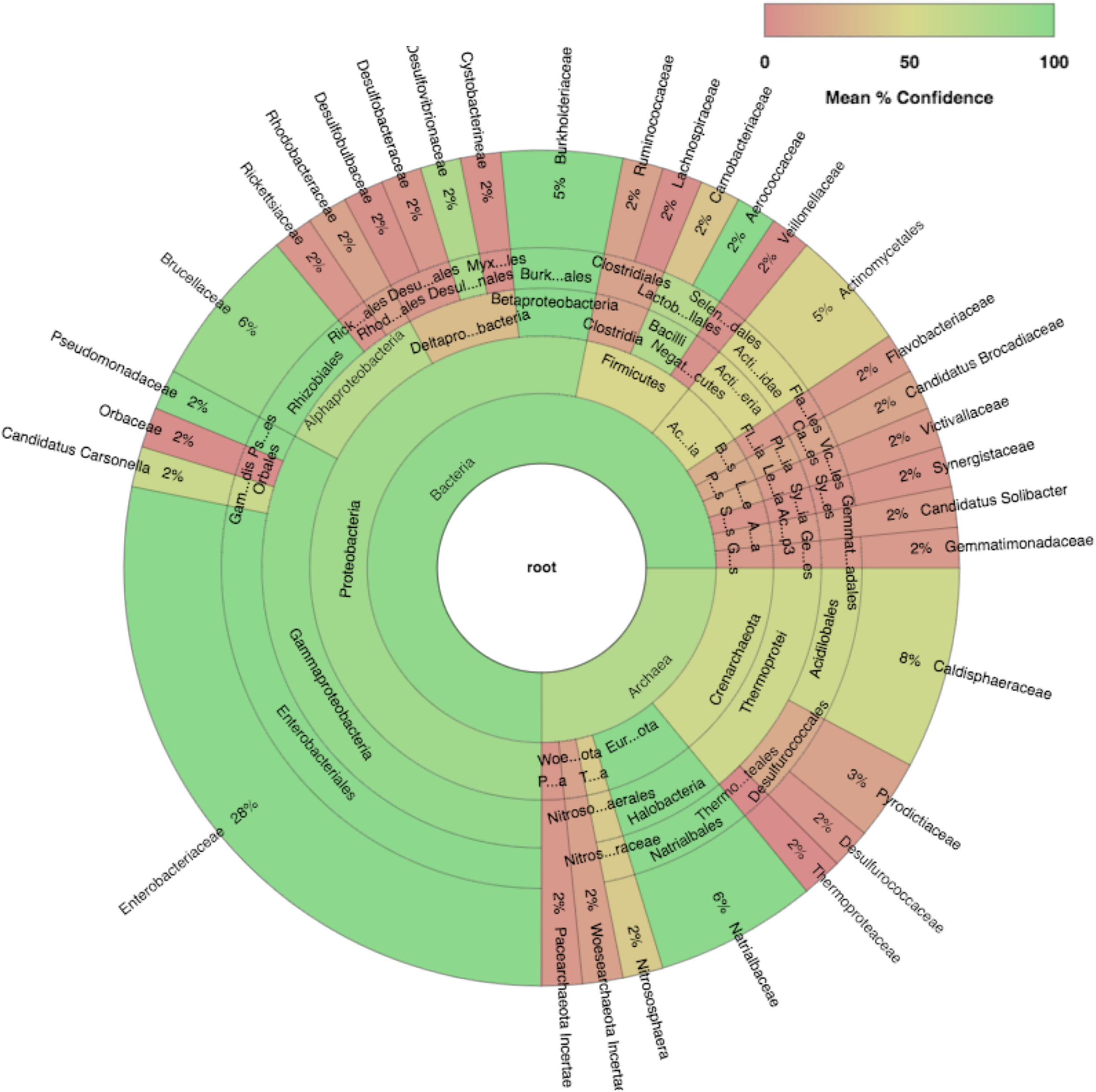
Microbial community profile identified in the intestine of *A. obliqua*. The 16S rDNA bacterial, actinomycete, and archaea clones obtained from wild and laboratory adult and larvae flies were curated, merged and analyzed in Geneious software to determine 16S biodiversity. Most of the recovered sequences (~85%) were relative to bacteria showing three main clades (proteobacteria, firmicutes and actinomycetes) ~25% were relative to Archaea.

In this work, we present the inter-kingdom diversity harbored in the midgut of *A. obliqua*, expanding the intricate microbial crosstalk, which could be related to different biological outcomes (e.g., bio-assimilation) in larvae and adult wild or laboratory flies. It is important to note that the techniques based on the analysis of rRNA genes amplified by PCR may not represent a complete and accurate picture of the microbial community. The estimation of genetic diversity by DGGE is limited to members of dominant microbial communities, which must represent at least 1% of the total microbial population to produce a visible band in DGGE (Muyzer, de Waal and Uitterlinden, 1993). This study is only the beginning of our understanding of the intestinal ecosystem of *Anastrepha*. We have determined some predominant populations, which could significantly modulate the fitness and physiology in mass-rearing and wild *A. obliqua* flies through an inter-kingdom communication of such communities. Further studies are required to understand how these microorganisms affect metabolic processes, sexual behavior, and other aspects of the fly’s physiology. This knowledge will eventually improve the mass rearing technology of *A. obliqua*, and focus on the discovery of microorganisms and enzymes of nutritional interest for fruit flies.

## Supporting information

Electronic Supplementary Material

## Acknowledgements

This research was financed by the National Council of Science and Technology project CB-2008-01-101389, and fellowship to GZ-R number 17171.

## Electronic Supplementary Material

**Online Resource 1** Table - NCBI accession number of the sequences used in this study.

**Online Resource 2** DGGE analysis of 16S biodiversity of bacteria, archaea, and fungi communities associated at the intestine of A. obliqua.

**Online Resource 3** Electropherogram of 16S rDNA clones.

**Online Resource 4** 16S biodiversity of Bacteria and Actinomycete communities associated at the intestine of A. obliqua based on 16S rDNA sequences.

**Online Resource 5** 16S biodiversity of Archaea communities associated at the intestine of A. obliqua based on 16S rDNA sequences.

