## Supplementary material for "The fruit fly *Anastrepha obliqua* harbors three kingdoms of life in its intestinal tract": Electronic Supplementary Material

Title:

El Colegio de la Frontera Sur, Carretera Antiguo Aeropuerto Km. 2.5, Tapachula, Chiapas, Mexico.

[2] Programa Moscafrut, SENASICA-SAGARPA Camino a Cacaotales, S/N. C.P. 30860, Metapa de Domínguez, Chiapas, Mexico

**Online resource 1**

Table - NCBI accession number of the sequences used in this study.

| Submission code | Sample code | NCBI Number | Submission code | Sample code | NCBI Number |
|---|---|---|---|---|---|
| SUB5244878 | LA-B6 | **MK729487** | SUB5244878 | WL_Ar4 | **MK729511** |
| SUB5244878 | LA-B2 | **MK729488** | SUB5244878 | WL_Ar5 | **MK729512** |
| SUB5244878 | LA-B6_2 | **MK729489** | SUB5244878 | WL_Ar6 | **MK729513** |
| SUB5244878 | LA-B8 | **MK729490** | SUB5244878 | WL_Ar7 | **MK729514** |
| SUB5244878 | LL-A15 | **MK729491** | SUB5244878 | WL_Ar8 | **MK729515** |
| SUB5244878 | LL-B1 | **MK729492** | SUB5244878 | WL_Ar9 | **MK729516** |
| SUB5244878 | LL-B2 | **MK729493** | SUB5244878 | LL_Ar1 | **MK729517** |
| SUB5244878 | LL-B3 | **MK729494** | SUB5244878 | LL_Ar2 | **MK729518** |
| SUB5244878 | LL-B4 | **MK729495** | SUB5244878 | LL_Ar3 | **MK729519** |
| SUB5244878 | LL-B6 | **MK729496** | SUB5244878 | LL_Ar4 | **MK729520** |
| SUB5244878 | WA-B3 | **MK729497** | SUB5244878 | LL_Ar5 | **MK729521** |
| SUB5244878 | WA-A2 | **MK729498** | SUB5244878 | LL_Ar6 | **MK729522** |
| SUB5244878 | WL-Ac2 | **MK729499** | SUB5244878 | LL_Ar7 | **MK729523** |
| SUB5244878 | LA-B3 | **MK729500** | SUB5244878 | LL_Ar9 | **MK729524** |
| SUB5244878 | WL-B1 | **MK729501** | SUB5244878 | LL_Ar10 | **MK729525** |
| SUB5244878 | WL-B2 | **MK729502** | SUB5244878 | LA_Ar1 | **MK729526** |
| SUB5244878 | WL-B4 | **MK729503** | SUB5244878 | LA_Ar2 | **MK729527** |
| SUB5244878 | WL-B6 | **MK729504** | SUB5244878 | LA_Ar3 | **MK729528** |
| SUB5244878 | WA-Ar2 | **MK729505** | SUB5244878 | LA_Ar4 | **MK729529** |
| SUB5244878 | WA-Ar8 | **MK729506** | SUB5244878 | LA_Ar5 | **MK729530** |
| SUB5244878 | WA-Ar9 | **MK729507** | SUB5244878 | LA_Ar8 | **MK729531** |
| SUB5244878 | WA-Ar13 | **MK729508** | SUB5244878 | LA_Ar9 | **MK729532** |
| SUB5244878 | WA-Ar14 | **MK729509** | SUB5244878 | LA_Ar10 | **MK729533** |
| SUB5244878 | WA_q27 | **MK729510** | | | |

Submission code represents a code given during submission at NCBI. Sample code: Wild Adult (WA), Wild Larvae (WL); and Laboratory Adult (LA), and Laboratory Larvae (LL). B for bacteria, A or Ac for actinomycetes, Ar or q for archaea origin. NCBI number represent the accession number

**Online resource 2** DGGE analysis of 16S biodiversity of bacteria, archaea, and fungi communities associated at the intestine of A. obliqua. Bisacrylamide gel showing the bands obtained from the DGGE technique on amplification of 16S sequences of bacterial and Gram-positive enriched population (A & A'), Archaea (B), and Fungi (C) are shown. Fly samples are tagged as Wild Adult (WA), Wild Larvae (WL); and Laboratory Adult (LA), and laboratory Larvae (LL), FL refers to the food leg of fruit fly which was employed as a control of 18S amplicons.

**Online resource 3** Electropherogram of 16S rDNA clones.  Electropherograms of the first 600 nucleotides of one NCBI's accepted bacterial sequence LA_B6 (A) and its comparison with NCBI filtered sequences of Gram-positive bacteria WL_A5 and archaea WA_q4 (B & C, respectively) obtained in this study are shown. Sequences that showed good quality such as NCBI accepted sequences and depicted here were manually analyzed, trimmed, and subsequently analyzed to 16S diversity assay on Geneious.

**Online resource  4** 16S biodiversity of Bacteria and Actinomycete communities associated at the intestine of A. obliqua based on 16S rDNA sequences. The 16S rDNA bacterial and Gram-positive enriched clones obtained from wild and laboratory adult and larvae flies were merged and analyzed in Geneious software to 16S biodiversity. Sequences identity were relative in 89% to bacteria, 8% to actinomycete, and 3% to archaea.

**Online resource 5** 16S biodiversity of Archaea communities associated at the intestine of A. obliqua based on 16S rDNA sequences. The 16S rDNA archeae clones obtained from wild and laboratory adult and larvae flies were merged and analyzed in Geneious software to 16S biodiversity. ~56% of sequences were relative to archaea and ~45% to bacteria.

61 **Online resource 2**

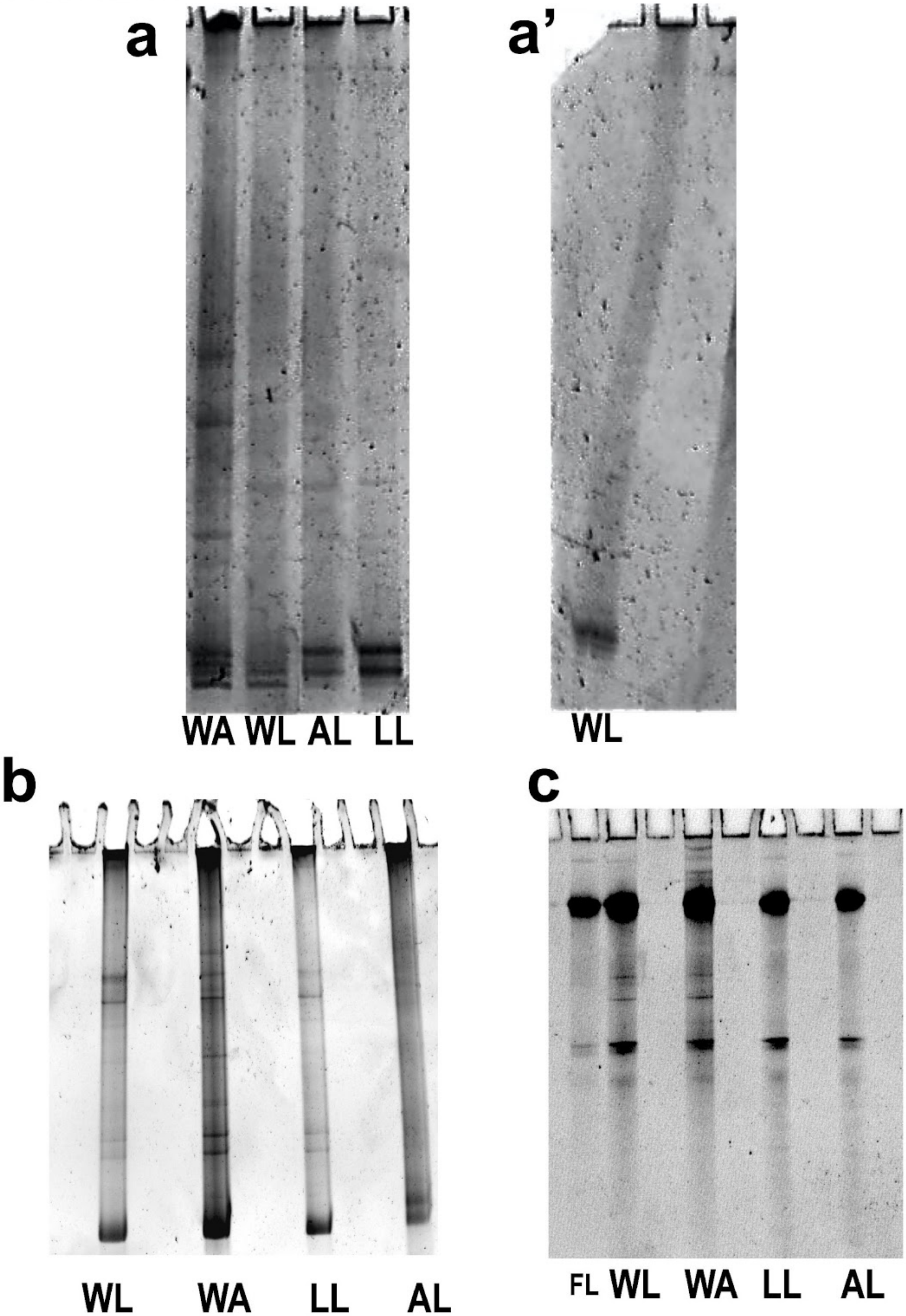

**Online resource 3**

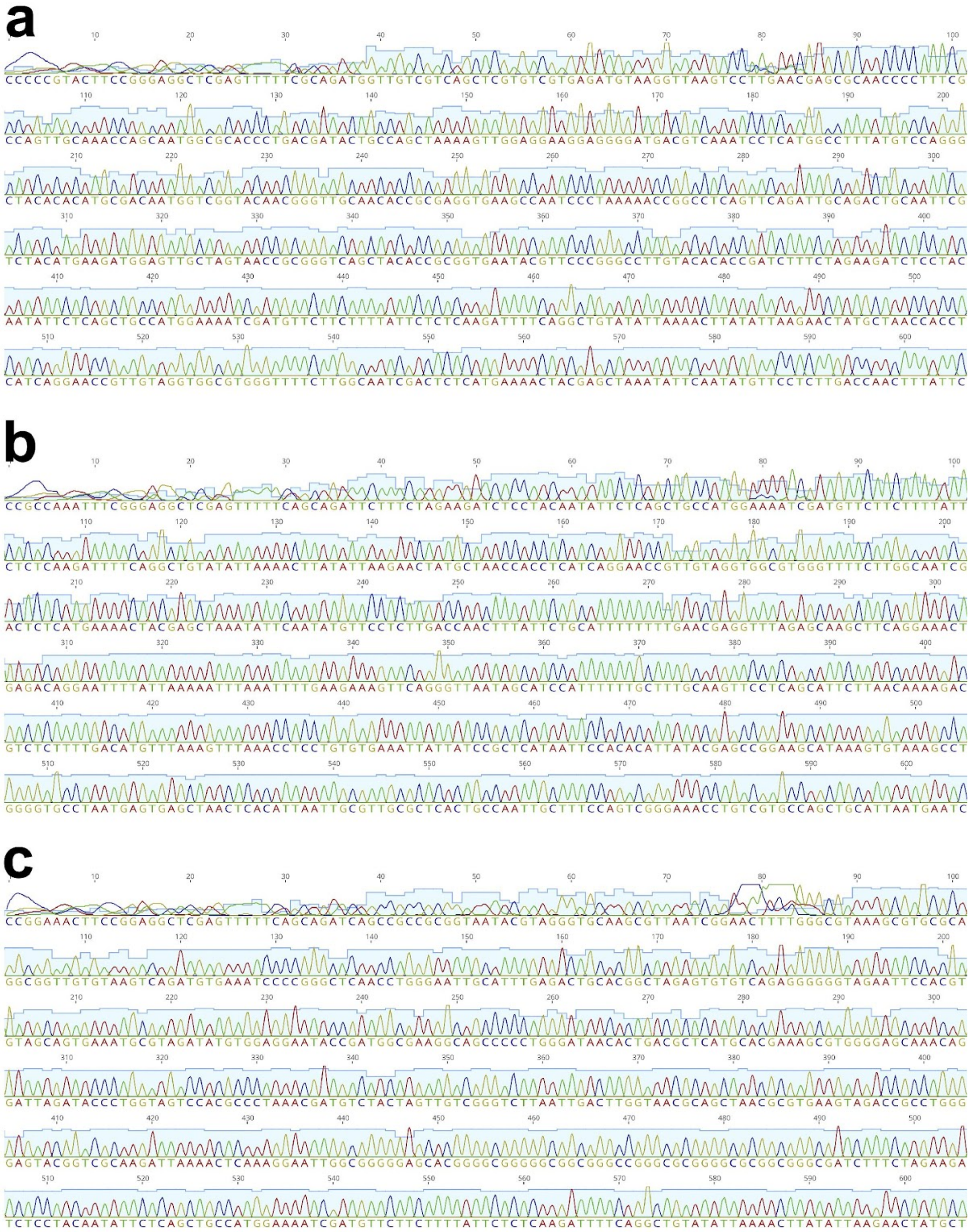

**Online resource 4**

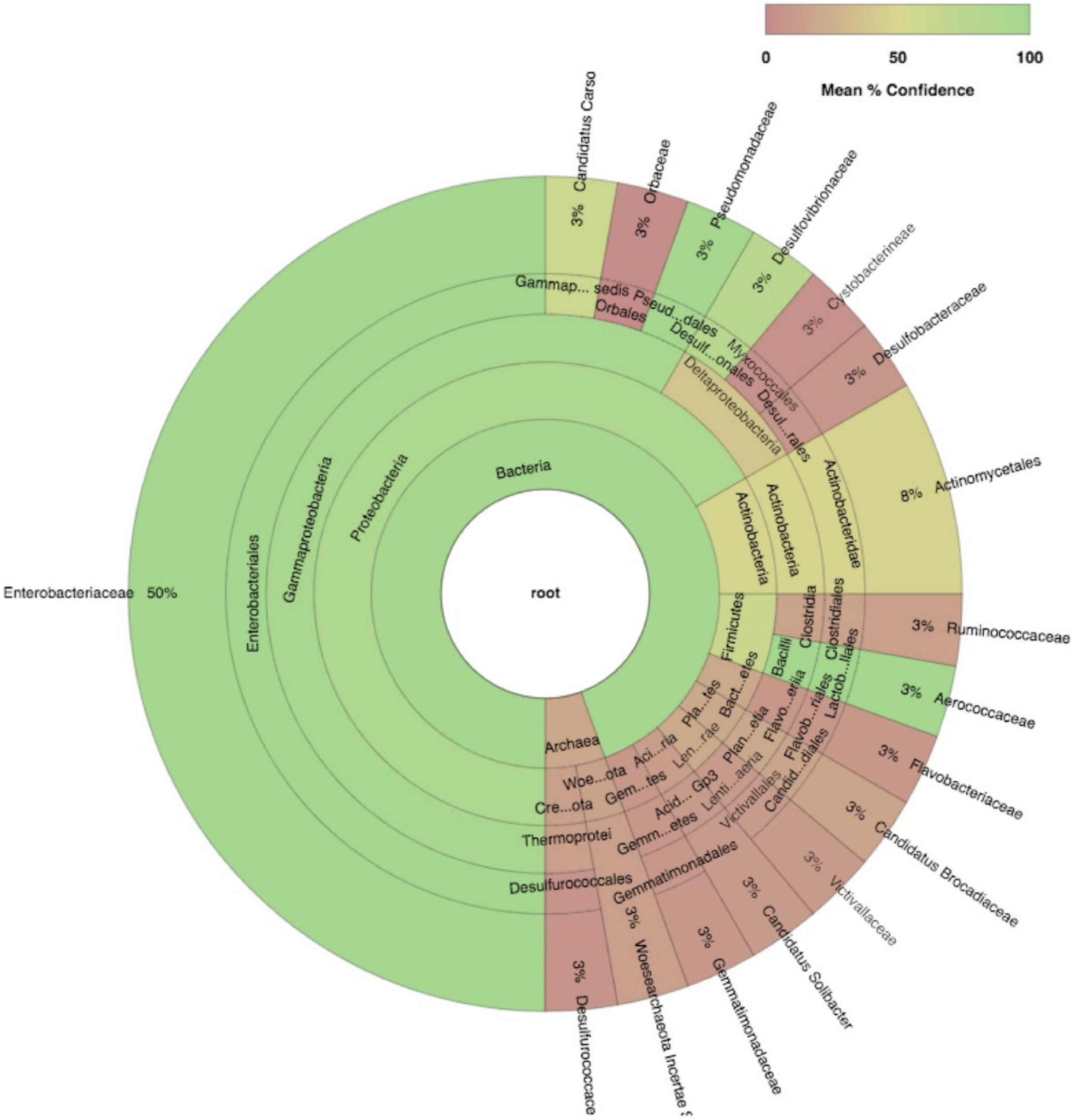

**Online resource 5**

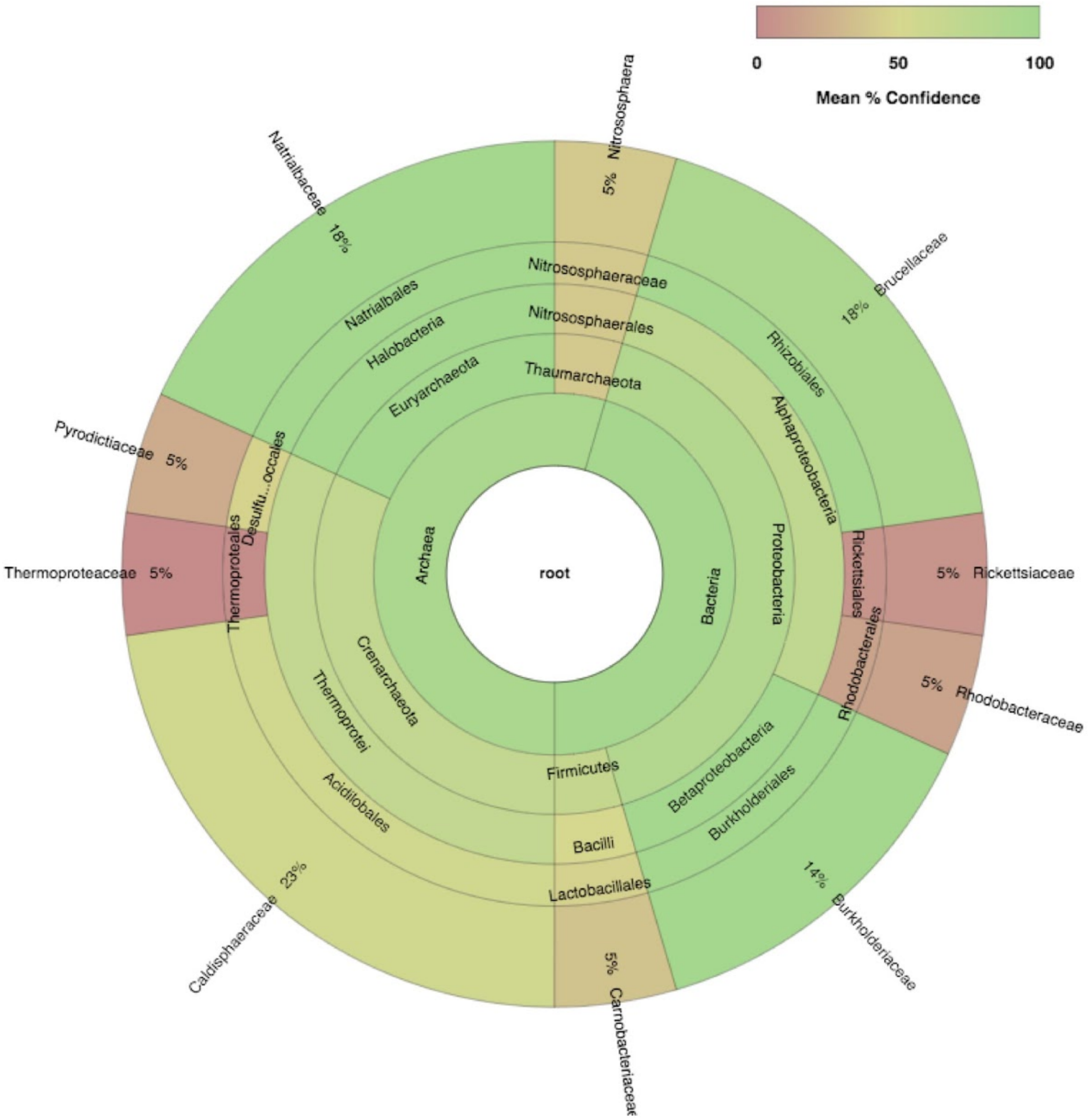
